# New genomic data and analyses challenge the traditional vision of animal epithelium evolution

**DOI:** 10.1101/228452

**Authors:** Hassiba Belahbib, Emmanuelle Renard, Sébastien Santini, Cyril Jourda, Jean-Michel Claverie, Carole Borchiellini, André Le Bivic

**Affiliations:** Structural and Genomic Information Laboratory, Aix-Marseille Université & CNRS UMR 7256, Mediterranean Institute of Microbiology (IMM FR 3479), Marseille, France.; Aix Marseille Univ, Univ Avignon, CNRS, IRD, UMR 7263, Mediterranean Institute of Marine and Continental Biodiversity and Ecology (IMBE), Station Marine d'Endoume, Marseille, France.; Aix-Marseille University, CNRS, UMR 7288, Developmental Biology Institute of Marseille Luminy (IBDM) Marseille, France.

**Keywords:** Epithelium evolution, non-bilaterian animals, cell polarity, cell-cell junctions

## Abstract

The emergence of epithelia was the foundation of metazoan expansion. To investigate the early evolution of animal epithelia, we sequenced the genome and transcriptomes of two new sponge species to characterize epithelial markers such as the E-cadherin complex and the polarity complexes for all classes (Calcarea, Demospongiae, Hexactinellida, Homoscleromorpha) of sponges (phylum Porifera) and compare them with their homologs in Placozoa and in Ctenophora. We found that Placozoa and most sponges possess orthologs of all essential genes encoding proteins characteristic of bilaterian epithelial cells, as well as their conserved interaction domains. In stark contrast, we found that ctenophores lack several major polarity complex components such as the Crumbs complex and Scribble. Furthermore, the E-cadherin ctenophore ortholog exhibits a divergent cytoplasmic domain making it unlikely to interact with its canonical cytoplasmic partners. These unexpected findings challenge the current evolutionary paradigm on the emergence of epithelia.

**SIGNIFICANT STATEMENT:** Epithelial tissues are a hallmark of metazoans deeply linked to the evolution of the complex morphogenesis processes characterizing their development. However, studies on the epithelial features of non-bilaterians are still sparse and it remains unclear whether the last common metazoan ancestor possessed a fully functional epithelial toolkit or if it was acquired later during metazoan evolution. In this work, we demonstrate that if sponges have a well conserved and functionally predicted epithelial toolkit, Ctenophores have either divergent adhesion complexes or lack essential polarity complexes. Altogether, our results raise a doubt on the homology of protein complexes and structures involved in cell polarity and adhesive type junctions between Ctenophora and Bilateria epithelia.

## Introduction

Multicellular organisms evolved from unicellular ancestors on several occasions during evolution of life ^1,2^ resulting in an extensive morphological diversity. For metazoans, this major transition is linked with the emergence of a new type of cellular organization, the epithelium^3–6^. Historically, epithelia were defined in bilaterians by the presence of three major features: apico-basal cell polarity, cell-cell junctions between the apical and the lateral domains and the presence of a basement membrane. These features are central to key epithelial functions: the regulation of internal/external exchanges and morphogenesis^6^. The typical bilaterian epithelium organization was extended by analogy to all eumetazoan i.e including Cnidaria and Ctenophora^7–9^ with very little molecular evidences preventing evolutionary interpretations of epithelial structures^4,9,10^.

From a morphological point of view, non bilaterian animals display a variety of cell sheet organization. For example, the basal lamina is absent from all but one sponge classes ^11,12^, in placozoa^13^ and in several ctenophoran species^14^. From a functional point of view, these epithelial-like cell layers show selective transport differences with bilaterian ones ^4,15–17–18^.It is now essential to determine the identity of the genes and proteins involved in these basal metazoan tissues – and consequently their homology across animals-remains to be determined ^5^.

Despite the diversity of epithelial structures among animals, apico-basal cell polarity and AJs are believed to be present in all extant animal phyla ^3,5,6,8,15,21^. We thus chose to characterize molecularly these two bilaterian epithelial hallmarks among non bilaterian phyla. Former studies performed on Placozoa^5,13,19,20,22^ and sponges^23–25^ initiated the study of candidate epithelial genes in the different lineages. The conservation of critical functions was not assessed, however, due to the lack of detailed analyses of key protein interaction domains and residues. On the other hand, studies on the epithelial organization of Ctenophora were neglected in favor of studies focused on the mesoderm and nervous system ^26–32^ due to the unquestioned position of this phylum among eumetazoans until recently ^33–35,^

In the present study, we first sequenced the genomes of two additional sponge species, *O. lobularis* (belonging to the Homoscleromorpha class) and *O. minuta* (the first Hexactinellida), and used RNA-seq data to help with the annotation procedure. This new data was then combined with information available from public databases to carefully identify and analyze homologs of genes coding for proteins known to compose polarity complexes and adherens junctions in all classes of Porifera (Calcarea, Demospongiae, Hexactinellida, Homoscleromorpha), several genera of Ctenophora with contrasted features ^14^ and Placozoa. Classical cadherins (E-type) and catenins ^5,36^, Par, Crumbs (Crb) apical polarity complexes and Scribble (SCRIB) lateral polarity complex were identified and analyzed ^8,37–40^. We hypothesize that sponge species exhibit highly contrasted tissue features related to molecular divergence of some of their polarity complex proteins. Finally, we revealed an unexpected lack of conservation of the epithelial toolkit in Ctenophores asking for a profound revision of our understanding of Ctenophore biology. Altogether, our results raise a doubt on the homology of protein complexes and structures involved in cell polarity and adhesive type junctions between Ctenophora and Bilateria epithelia.

## Results

### New genomic and transcriptomic data from *Oopsacas minuta* (Hexactinellida) and *Oscarella lobularis* (Homoscleromorpha)

We used two platforms (Pacific Bioscience and Illumina) and a combination of paired end and mate pair sequencing approaches (see Materials and Methods) to generate and assemble the data.

Concerning *Oopsacas minuta*, the assembly yielded a total of 61.46 Mb of unique haploid genome sequence distributed in 365 contigs longer than 1 kb (N_50_ length=676,369 bp, L_50_ number=31, mean coverage=381). Following the mapping of 207,529,788 RNAseq reads from a polyA+ cDNA library (mean Open Reading Frame (ORF) coverage = 1443), we predicted the presence of 17,043 protein-coding genes. The small final number of contigs and the above coverage values suggest that our delineation of the (protein-coding) gene content is very close to 100% completion.

Concerning *Oscarella lobularis*, we generated and assembled a total of 52.34 Mb of unique haploid genome sequence distributed in 2,658 contigs longer than 1 kb (N_50_ length=265,395 bp, L_50_ number=58, mean coverage=98). Following the mapping of 231,475,388 RNAseq reads from a polyA+ cDNA library (mean ORF coverage = 710), we predicted the presence of 17,885 protein-coding genes. The large, albeit unavoidable, proportion (>50%) of sequence data from bacterial and archaean origin, as well as unfavorable (repeated or variable) genome structures caused the final number of contigs to remain significantly larger than for *O. minuta.* However, the above coverage values remain large enough to correspond to a near 100% complete delineation of the (protein-coding) gene content.

The quality of our transcriptomes and genome drafts enables us to be confident on the completion of the predicted proteins and the number of copies found for each candidate gene.

### Porifera common ancestor most likely possessed functional adherens junctions

Classical cadherins contain extracellular repeat domains that mediate trans-interactions with the extracellular domain of cadherins on opposing cells, and a cytoplasmic domain that binds p120 and β-catenin ^5,41–43^. β-catenin binds to α-catenin thereby forming the core cytoplasmic protein complex of the classical cadherin/β-catenin/α-catenin complex (CCC). In this complex, α-catenin is the key protein that links the CCC complex to the underlying actin cytoskeleton. In turn, p120 is the critical actor for the surface stability of cadherin-catenin cell-cell adhesion by controlling cytoskeletal dynamics and regulating cadherin endocytosis.

Cadherin and catenin families are present outside of metazoans, but the C-terminal catenin-binding motifs that define classical E-cadherins are a metazoan novelty^5,36,44^. Consistent with earlier reports, our analyses, combining homology searches, phylogenetic reconstructions and domain predictions, confirm that Placozoa and all Porifera possess homologs of classicalE-cadherins ^5,13,19,29,22–25,36,42^. All characteristic domains were identified with high confidence (Fig. 1A):

- The extracellular cadherin (EC) repeated domains (ranging from 3 in *Sycon ciliatum* to 32 in *Trichoplax adhaerens* units) that mediate trans-interactions with the extracellular domain of cadherins on opposing cells;
- The transmembrane region (TM) and the cytoplasmic tail, which contains the conserved specific binding domain for p120-catenin in the juxta-membrane domain (JMD) and the β-catenin-binding domain (CBD);
- The epidermal growth factor (EGF) domains and Laminin G (Lam-G) domains in a membrane-proximal position considered typical of non-vertebrate classical cadherins ^21,42^.

**Figure 1.**
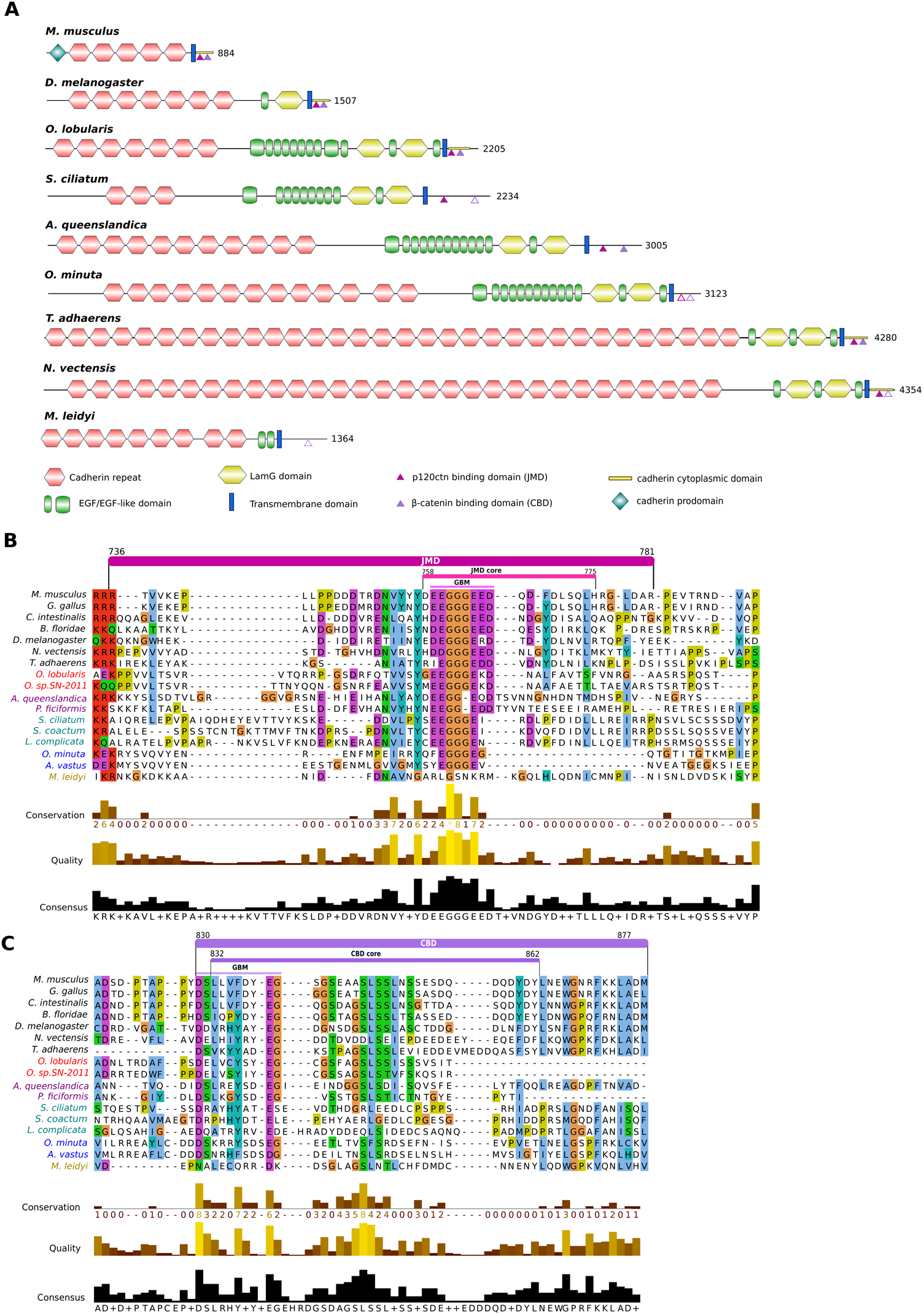
Comparison of E-cadherin domains and motifs between metazoans. Porifera: Homoscleromorpha in red (*Oscarella lobularis*, *Oscarella. sp.*), Demospongiae in magenta (*Amphimedon queenslandica*, *Petrosia ficiformis*), Calcarea in green (*Sycon ciliatum*, *Sycon coactum*, *Leucosolenia complicata*), Hexactinellida in blue (*Oopsacas minuta*, *Aphrocallistes vastus*). Other represented clades are Placozoa(*Trichoplax adhaerens*); Ctenophora (*Mnemiopsis leidyi*) in yellow, Cnidaria (*Nematostella vectensis*), Bilateria (Deuterostomia: *Mus musculus;* Protostomia: *Drosophila melanogaster*). Sequences were aligned with MAFFT v7 web server and visualized with Jalview. **A) Representative cadherin proteins depicted with their domains**. *Mus musculus* and *Drosophila melanogaster* E-cadherins are taken as reference. *Oscarella lobularis* has the sole poriferan cadherin the cytoplasmic-specific domain of which is detected by Pfam (E-value=2.10^-11^) and InterProScan as in the mouse and fruitfly E-cadherin (depicted in yellow at the C-terminal part). Degrees of conservation of p120 and β-catenin binding domains are indicated by full, dashed or open triangles. **B) Alignment of the cytoplasmic cadherin p120 binding domain (Juxtamembrane domain, JMD).** The JMD consists of 50 residues immediately following the transmembrane domain (in Mouse E-cadherin). The JMD core consists of 20 residues. The groove-binding motif (GBM) required for binding p120 is well conserved in metazoans. **C) Alignment of the cytoplasmic cadherin β-catenin binding domain (CBD).** The CBD consists of approximately 50 residues. The groove-binding motif (GMB) consists of 10 residues.

The alignment of E-cad JMD that mediates binding to p120 catenin (Fig. 1B) shows that the Groove-Binding Motif (GBM motif) (XX[ED]GGGEXX) is highly conserved in placozoan and in three classes of sponges. In contrast a G residue is missing in the two demosponges studied, which may modify the interactions with p120-catenin, since the three consecutive glycine residues are thought to anchor the region in a small hydrophobic pocket in the armadillo (ARM) repeats of p120-catenin ^42,45^. p120-catenin consists of central ARM domain repeats involved in E-cad JMD interactions flanked by an N-terminal regulatory region (NTR) and a C-terminal tail region (CTR). Among the key residues of p120 involved in E-cadherin-binding (Fig.S1A), the 13 essential residues involved in electrostatic interactions (Q391 to K574) with the N-terminal acidic region of the JMD core (residues758-766, ^45^(Fig. 1B) are highly conserved in sponges and placozoans with minor exceptions (H392 ->Q and K574->Q residues) in glass (Hexactinellid) sponges. In contrast, eight amino acids in the N-terminus of p120 (R364 to Y389, Figure S1A) known to be involved in hydrophobic interactions with the C-terminal anchor region of the JMD core (residues767-775) are more variable. This region (Fig. 1B) of E-cadherin appears also less constrained suggesting that electrostatic interactions dominate the p120-catenin-JMD core interaction.

We detected a striking exception in the ctenophore *Mnemiopsis leidyi*, in which the E-cadherin GBM motif is not conserved (Fig. 1B) possibly forbidding interaction with p120-catenin. In contrast, *M. leidyi* p120-catenin residues, essential for electrostatic binding with E-cadherin classical cytoplasmic domain, remain highly conserved (Fig.S1A), thus excluding a compensatory co-evolution process ^44^that may have preserved the interaction. In the cadherin domain binding to β-catenin (CBD, Fig. 1C), the interaction was shown to require a GBM of about 10 amino acids (DXXXXϕXXEG where ϕ is an aromatic residue) ^42,45,46^. As for the p120-catenin-binding motif, this motif is conserved in *Trichoplax* and all sponges except in calcareous sponges where it slightly diverges at the end (Fig. 1C). Whether such a change in the CBD results in a weakening (or loss) of the interaction with β-catenin in calcareous sponges has yet to be investigated.

A single β-catenin gene copy was identified in each studied species except for calcareous sponges that exhibit a specific duplication (Fig. S1B). All β-catenin proteins identified in sponges, placozoans and ctenophores harbor the same ARM repeats as described in bilaterians. In sponges and placozoans, β-catenin residues that were identified as essential for E-cadherin interaction ^47^are highly conserved except for the R386 and N387 residues (respectively replaced by L and T) in two hexactinellids and a more anecdotic change: A656 -> S in placozoans.

In *M. leidyi* again the E-cadherin CBD motif diverged from that of other metazoans. The D, the aromatic residue, and the G were replaced in the DXXXXϕXXEG sequence (Fig. 1C), which might impair interactions with the classical β-catenin respective interaction motif. Conversely *M. leidyi* β-catenin amino acids involved in the interaction with E-cadherin (Fig. S1B) are modified (Y331V, K335Y, D390N and R582C) either suggesting a loss of interaction with the E-cadherin CBD intracellular domain or its maintenance through co-evolution.

The α-catenin (member of the vinculin family) links E-cadherin to the actin cytoskeleton by interacting with β-catenin ^36^. All species studied, including *T. adhaerens* and *M. leidyi*, have at least two vinculin family members: one orthologous to a-catenin and one orthologous to vinculin. Interestingly, the a-catenin/vinculin N-terminal region, known to interact with β-catenin, is conserved (Fig. S1C). In addition, in the β-catenin of sponges, placozoans and ctenophores, most of the crucial residues involved in a-catenin binding ^47,48^are conserved (Fig.S1B) suggesting that such an interaction was already present in the common ancestor of metazoans. Our analyses of two additional sponge species (*Oscarella lobularis*, class Homoscleromorpha; *Oopsacas minuta*, class Hexactinellida) confirm the presence of *bona fide* E-cadherin complexes in the four Porifera classes. Moreover, the motifs governing the interactions between the members of this CCC complex essential for the establishment of adherens junctions appear very conserved in Placozoa, Calcispongiae and Homoscleromorpha. Even if a few substitutions were identified in demosponges and glass sponges that may modulate these interactions and explain the absence of AJs in these two lineages, we can nevertheless infer that the last ancestor of Porifera already possessed all the component needed to build functional adherens junctions similar to those of bilaterians. However, this is probably not true of the ctenophores as their E-cadherin cytoplasmic domain lacks most *bona fide* E-cadherin cytoplasmic domain binding sites.

### A Par apical polarity complex inherited from Urmetazoa

Next we investigated whether placozoans, ctenophores, and sponges of all classes, harbor the polarity protein complexes that are necessary for epithelium formation and morphogenesis ^37^. There are at least three types of polarity complexes:

- The Par complex made of atypical Protein Kinase C (aPKC), Partition defective 3 (Par3) and Partition defective 6 (Par6);
- The Scribble complex made of Scribble (Scrib/Src), lethal giant larvae (Lgl) and Disc large (Dlg);
- The Crumbs complex made of Crumbs (Crb), stardust (Sdt, or MPP5 (Membrane Palmitoylated Protein 5) in mammals also known as Pals1 (protein associated with Lin-7 1) and Pals1-associated tight junction protein (Patj).

First, we looked for the Par complex, considered to be a metazoan innovation ^23^. Par6 contains an N-terminal Phox and Bem1 (PB1) domain, a C-terminal Postsynaptic-density-95/Disc-large/Zona-occludens1 (PDZ) domain and a semi-CRIB (Cdc42/Rac interactive binding domain) motif immediately preceding the PDZ domain. The PB1 domain of Par6 forms a heterodimer with the PB1 domain of aPKC. Par3 is associated with Par6/aPKC complex via the PDZ-PDZ domain interaction. The activity of the PAR complex is dynamically regulated by phosphorylation of PAR3 and its association with the stable PAR6-aPKC complex. Highly conserved sequences for all members of this complex are present in all available genomes of sponges, placozoans and ctenophores. All characteristic domains as well as the residues essential for their interactions within the complex were also identified (Fig. 2, 3, S2).For example, Par6 (Fig. 2) interacts with aPKC through its PB1 domain and, in mouse Par6, lysine K19 is essential for this interaction ^49^. This Lysine residue is strictly conserved in all species studied here, which strongly suggests that the interaction between aPKC and Par6 is conserved throughout metazoan evolution. In addition, there are increasing evidences that the formation of this complex is regulated by phosphorylation, mainly on serine S980 in the aPKC-binding region of Par3 ^50^. This phosphorylated site (S/T) is conserved in all studied species (Fig. S2). All these data strongly suggest that aPKC, Par3 and Par6 have co-evolved from a functional metazoan ancestral complex.

**Figure 2.**
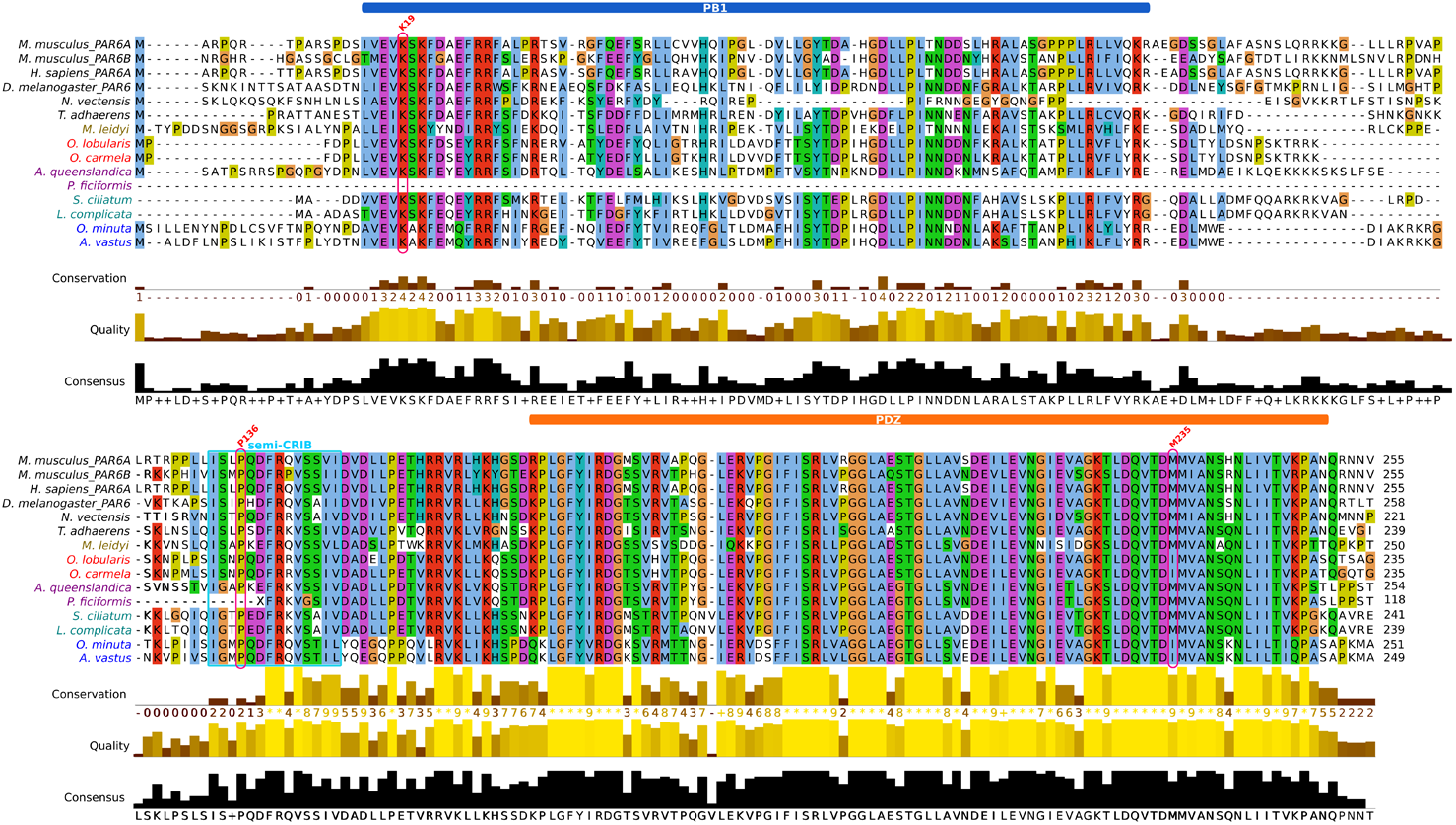
Comparison of the sequences of the diagnostic domains of Par6. The N-terminal Phox and Bem1 (PB1) domains required for interaction with the PB1 domain of atypical Protein Kinase C (aPKC); the semi-CRIB (Cdc42/Rac interactive binding) domain and the C-terminal Postsynaptic-density-95/Disc-large/Zona-occludens1 (PDZ) domain required for the interaction with Par3. Sequences were aligned with MAFFT v.7 and visualized with Jalview 2.9. Critical residues are labelled in red: Lysine K19 in PB1 domain is essential for the interaction with aPKC; Proline 136 (P136) in the semi-CRIB motif is necessary to bind cdc42; Methionine 235 (M235) in the PDZ domain binds the LGL protein.

**Figure 3.**
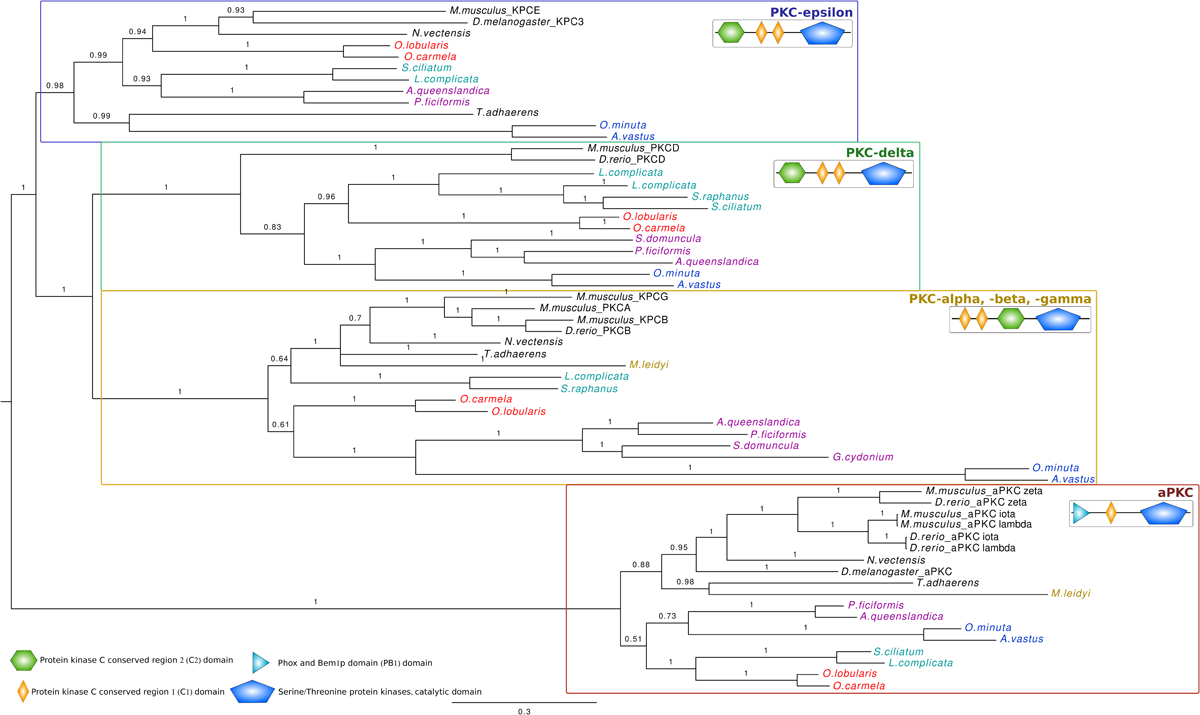
Phylogenetic relationships between members of the Protein Kinase C (PKC) family. A bayesian tree was inferred with available bilaterian sequences and predicted cnidarian, poriferans, placozoan and ctenophoran sequences aligned with MAFFT v7.123b. MrBayes was run under LG+G model of evolution with 4 rate categories and 1 million generations sampled every 1000 generations. The tree was rooted at midpoint and posterior probabilities are indicated for each branch. The canonical domain architecture was depicted for each PKC type. All non-bilaterian species studied have one copy of atypical Protein Kinase C (aPKC) according to both their domain composition and the robustness of the orthology group (pp=1).

### The Scribble lateral polarity complex is present in all non bilaterians except ctenophores

Next, we investigated the presence of the Scribble polarity complex composed of Scrib, Dlg and Lgl members. Members of this complex contain multiple protein-protein interaction domains, in particular PDZ, Src homology 3 (SH3) domain and guanylate kinase (GUK) domains capable of recruiting a complex network of proteins.

We identified with high confidence Dlg orthologous genes containing all specific domains (Lin2 and Lin7 binding domain (L27), GUK, SH3 and three PDZ domains) (Fig. 4) in all sponges and in *T. adhaerens* (even though the L27 domain is lacking). In ctenophores, Dlg orthologous genes were found without GUK domain. Since Dlg predates the emergence of metazoans (Dlg homologs have been reported in Choanoflagellata, Filasterea and Ichthyosporea)^23,36,51^ the absence of this key domain is probably due to secondary lost. Lgl, characterized by short WD40 repeats and specific phosphorylation sites, is not present in Choanozoa ^24^but was identified in all Porifera and Placozoa in agreement with previous studies ^23,24^and in Ctenophora (Table S1), suggesting that it appeared in the last common metazoan ancestor.

**Figure 4.**
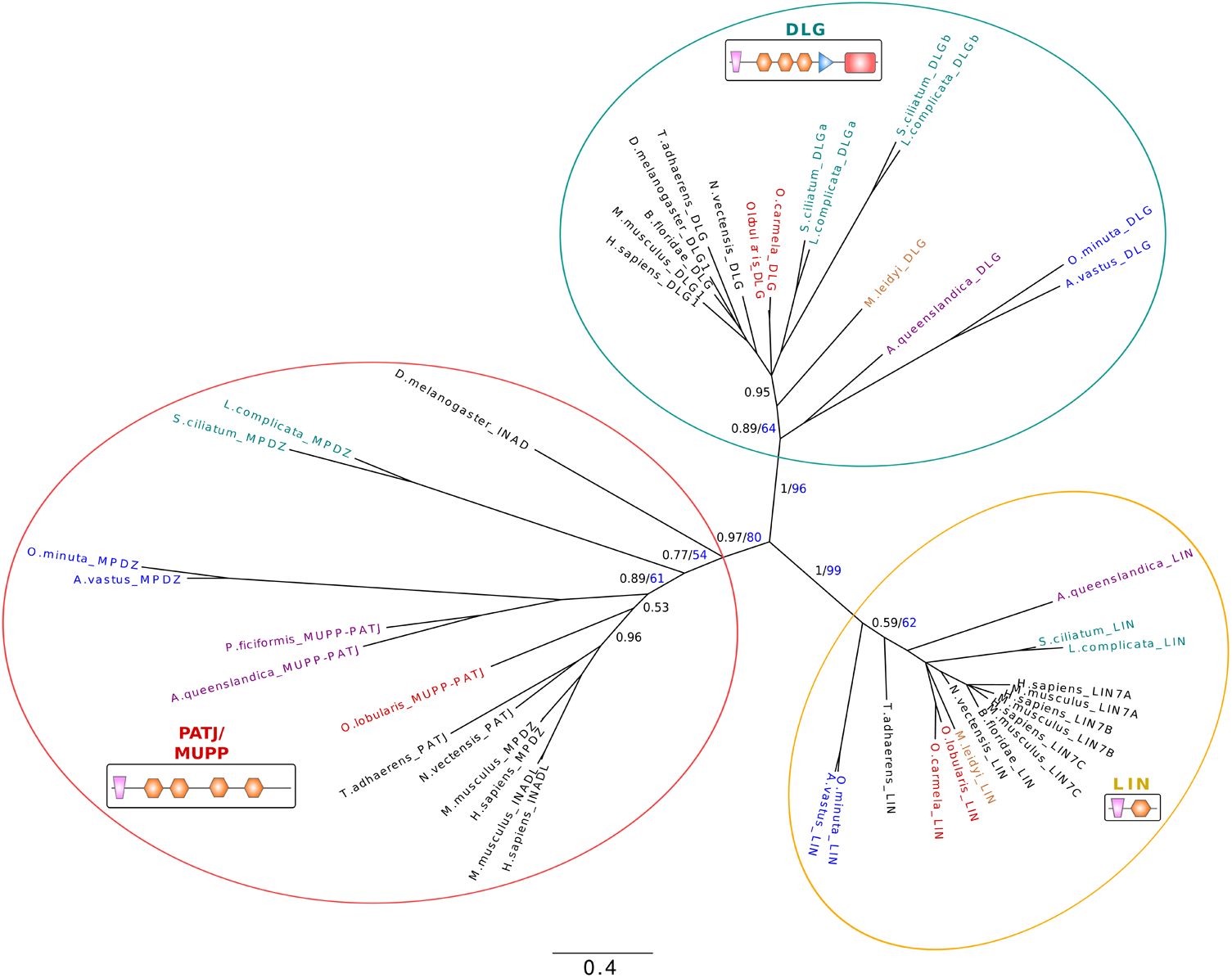
Phylogenetic relationships between GUK proteins. (Pat J, Lin and GLG proteins) based on their L27 and two first PDZ domain (except for LIN proteins which have a single PDZ) sequences. Available bilaterian sequences and predicted cnidarian, poriferan, placozoan and ctenophoran sequences were aligned with MAFFT v7.123b. The consensus phylogenetic tree was computed with PhyML and MrBayes. Both analyses were run under a LG evolution model with a gamma distribution and 4 rate categories. A total of 1 million generations, sampled every 1000 generations with a burn-in of 250 was used for the bayesian analysis. Bayesian posterior probabilities are shown in black and 100-bootstap PhyML replicates are shown in blue for each branch. Low-scoring L27 domains were also included in the alignment. Canonical domain architecture is depicted for PATJ-MUPP1, LIN and DLG family protein. We identified with high confidence Dlg orthologous genes coding for specific domains (Lin2 and Lin7 binding domain (L27), GUK, SH3 and three PDZ domains) in all sponges and in *T. adhaerens* (even though the L27 domain is missing). In ctenophores, Dlg orthologous genes were found without GUK domain. In Porifera, we found that all species possess PatJ homologs that cluster with bilaterian PatJ. In contrast, no PatJ homolog was found in ctenophores.

As previously reported, Scrib homologs were identified in all sponge classes ^24^and placozoans ^36^(Table S2). Scribble isa LAP [LRR (leucine-rich repeats) and PDZ (PSD-95/Discs-large/ZO-1) domain] protein containing16 LRRs and either one or four PDZ domains ^40,52^. In striking contrast, we could not detect a protein associating a PDZ domain and LRRs in ctenophores. To discard the hypothesis that divergent evolution led to a specific loss of Scribble in *M. leidyi*, we investigated its presence in two other genera (*Pleurobrachia* and *Beroe*) and we confirmed the absence of Scribble homolog (the only LRRs domains we identified belong to other classes). The LRR domain is critical for Scribble function, since in *Drosophila* Scribble proteins mutated in the LRR domains mimic the complete loss of Scribble protein^40^. The PDZ domain of Scribble was also shown to be important for its recruitment to the junctional complex and plasma membrane ^53^and for the correct localization of Dlg ^40^in *Drosophila*. The absence of a *bona fide* Scribble homolog in ctenophores might indicate a change in Dlg/Lgl localization or function.

### The Crumbs apical polarity complex is divergent in syncitial glass sponges and absent in ctenophores

We then investigated the conservation of the Crumbs complex in metazoans. Crumbs is a central regulator of epithelial apical actin cytoskeleton organization and adherens junction formation in bilaterians and was proposed to be a metazoan innovation ^8,23^ The formation of the Crumbs complex is ensured by physical interactions between different core components. The central component, Stardust/MPP5, organizes a plasma membrane-associated protein scaffold via an interaction between its PDZ domain and the C-terminal ERLI motif of Crb. The two L27 domains of Sdt bind to the L27 domains of PATJ and Lin-7.

Crumbs transmembrane proteins, consist of extracellular EGF, laminin-like repeats and a short cytoplasmic domain (less than 40 amino acids) with two essential sequence motifs ^37^(Fig. 5A). These motifs are the signature of Crumbs proteins and are essential for their morphogenetic function ^54,55^. The membrane proximal motif RxxxGxYxPSor FERM-binding motif (FBM) is required for the interaction with proteins of the ERM (Ezrin-Radixin-Moesin) family that associates with the actin cytoskeleton ^56^(Fig. 5B). The second motif consists of the last 4 amino acids, ERLI, at the C-terminus (Fig. 5B). It is a class II PDZ-binding motif (PBM), which interacts with stardust (MPP5)^57^and Par6 ^58^. There is a strong conservation of the class II PDZ-binding site with conservative variations (E/D-R/K-L/I-I/L) in three of the four sponge classes and in *Trichoplax*, most likely under evolutionary pressures maintaining the interaction with PDZ containing proteins (Sdt or Par6). A unique exception was found in hexactinellids were the ERLI motif in replaced by E**<u>T</u>**LI (Fig. 5B). This change allows the binding of class I PDZ domains instead of class II. Analysis of FBM in *Trichoplax* and Porifera reveals that hexactinellids exhibit the most divergent sequences with only two conserved residues (XxxxXxYxPX) while *O. lobularis* has a conserved FBM (RxxxGxYxPT) (Fig. 5B) suggesting that homoscleromorphs have a truly functional Crumbs complex while it might be defective in hexactinellids. Therefore, there might be a relationship between the loss of some protein interactions and the syncytial organization characteristic of this sponge lineage. In Calcarea and in Demospongiae, 4 and 3 of the 5 FBM residues are conserved. Placozoans also exhibit a conserved FBM with RxxxGxFxPS in one of their two Crumbs homologs (Fig. 5B). Another feature of the FBM in bilaterians is the presence of two phosphorylation sites recognized by aPKC (**T**xG**T**Yx), which regulates the binding to Moesin, an ERM protein ^59^. These two phosphorylation sites are absent from all sponge species and from *Trichoplax*, suggesting that this regulation by aPKC is an innovation shared by cnidarians and bilaterians (except for *Caenorhabditis elegans*) since at least one phosphorylation site is present in a Crumbs isoform of cnidarians (*Nematostella*) (Fig. 5B). Thus, even though previous studies identified Crumbs-like proteins in all sponge classes, these studies did not verify the conservation of key functional motifs ^24^required for their functional interactions. Here, we show that the high divergence of key Crumbs residues in glass sponges is hardly compatible with the formation of a fully conserved complex (Fig. 5B).

**Figure 5.**
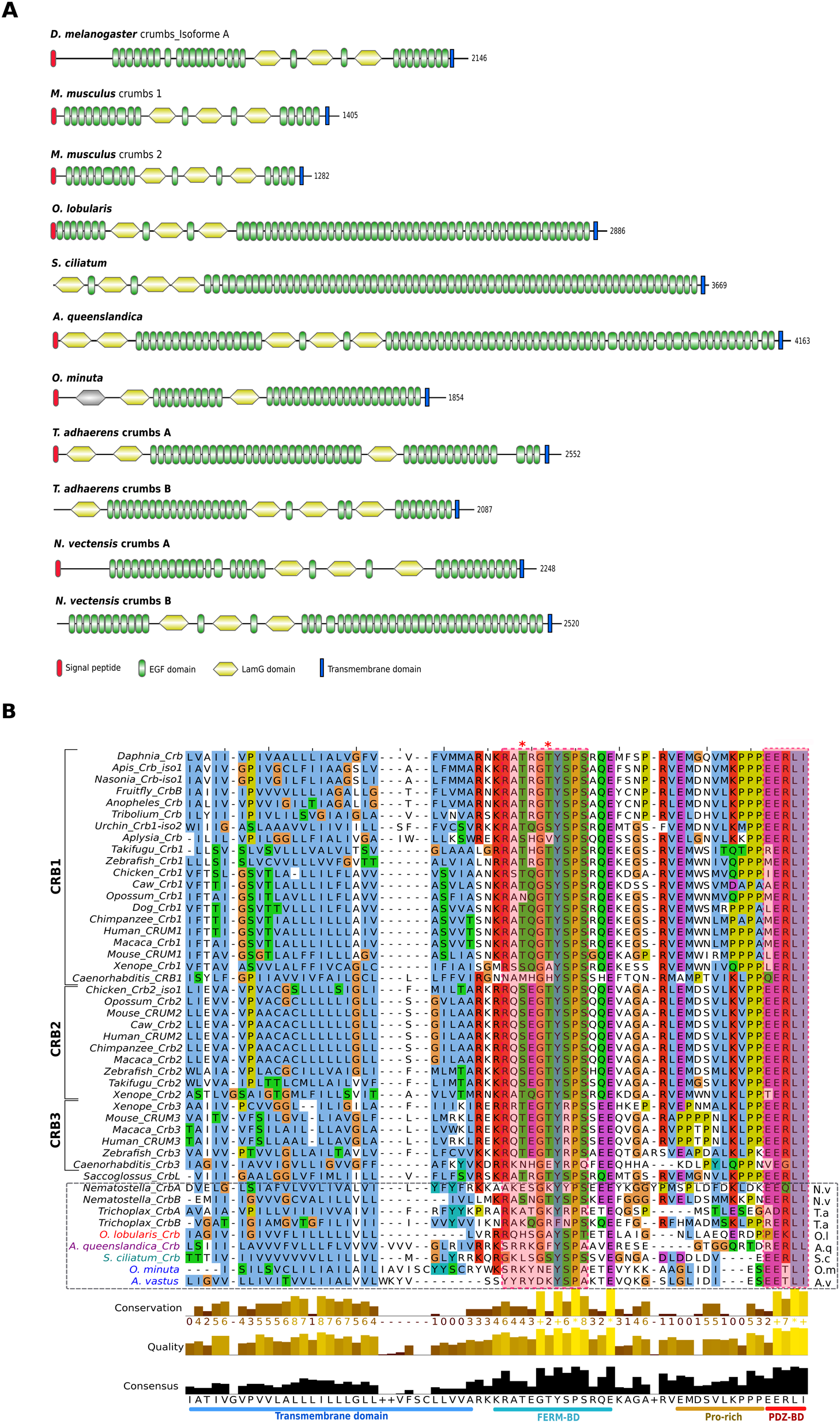
**A) Domain composition of Crumbs proteins** of *D. melanogaster* (Dcrb), *M. musculus* (CRB1, CRB2), *O. lobularis*, *S. ciliatum*, *A. queenslandica*, *O. minuta*, *N. vectensis* and *T. adhaerens*. Domains are shown as detected by SMART and Pfam and scaled with IBS software. Crumbs transmembrane proteins consist of extracellular epidermal growth factor (EGF), laminin-like (LAM) repeats and a short cytoplasmic domain. No crumbs was detected in Ctenophora. In contrast to other animals, sponges have only one copy of Crumbs. **B) Alignment of the cytoplasmic domain of Crumbs** that binds PALS1 and PAR6. Transmembrane and intracellular domains were aligned with MAFFT v7.123b and displayed with JalView. The transmembrane domain, the FERM binding domains (FBM) containing the RxxxGxYxPS motif needed for the interaction with the Ezrin-Radixin-Moesin (ERM) protein family, the Proline rich domain and the PDZ binding domain (PDZ-BD) are depicted at the bottom. The FBM presents a different conservation depending on the sponge class considered (from 2 to 5 conserved residues). Asterisks indicate the position of the two phosphorylation sites recognized by aPKC. These sites are absent in placozoans and poriferans. The PDZ-BD domain (interacting with MPP5) is well-conserved in non bilaterians except in glass sponges.

Finally, the most unexpected result was the absence of any Crumbs-like gene or transcript in *M. leidyi* (and in other ctenophore transcriptomes available in databases: *Pleurobrachia bachei*, *Beroe abyssicola and Beroe sp.*) exhibiting a significant similarity with the conserved cytoplasmic domain, while we identified homologs of transmembrane proteins with extracellular domains made of EGF and laminin-like repeats. This suggests that Crumbs proteins with a classical intracellular domain are present in all extant metazoans except ctenophores. Crumbs proteins interact with Sdt (MPP5 or Pals1 in mammals). Sdt encodes a membrane-associated guanylate kinase (MAGUK) protein containing two L27 domains, a single PDZ domain, a SH3 motif, a hook domain and a GUK domain ^37^. We identified orthologues of Sdt in all sponges as well as in Placozoa based on phylogenetic reconstructions (Fig. 6). However, we noticed that the first L27 domain by which Sdt is known to interact with the L27 domain of PatJ is absent in the two hexactinellid sponges and in Placozoa. In all cases, we could not detect MMP5 homologs in ctenophores despite the fact that other MPP genes or transcripts were presents (Fig. 6). The third partner of Crumb complex is the multiple PDZ domain containing protein PatJ which binds MPP5 via L27 interactions (Fig. S5). In Porifera, we found that all species possess PatJ homologs that cluster with bilaterian PatJ (Fig. 4), in contrast with a previous claim by Riesgo et al. (2014). However, we could not detect a L27 domain in hexactinellids (Fig. S5), which suggests a lack of interaction with Sdt/MPP5 proteins in glass sponges. In contrast, the characteristic L27 domain is present and highly conserved in the homoscleromorph sponge *Oscarella lobularis* and to a lesser extent in calcareans and demosponges. As no PatJ homolog was found in ctenophores, we can safely conclude that the whole Crumbs/Sdt/Patj complex is entirely absent in this phylum.

**Figure 6.**
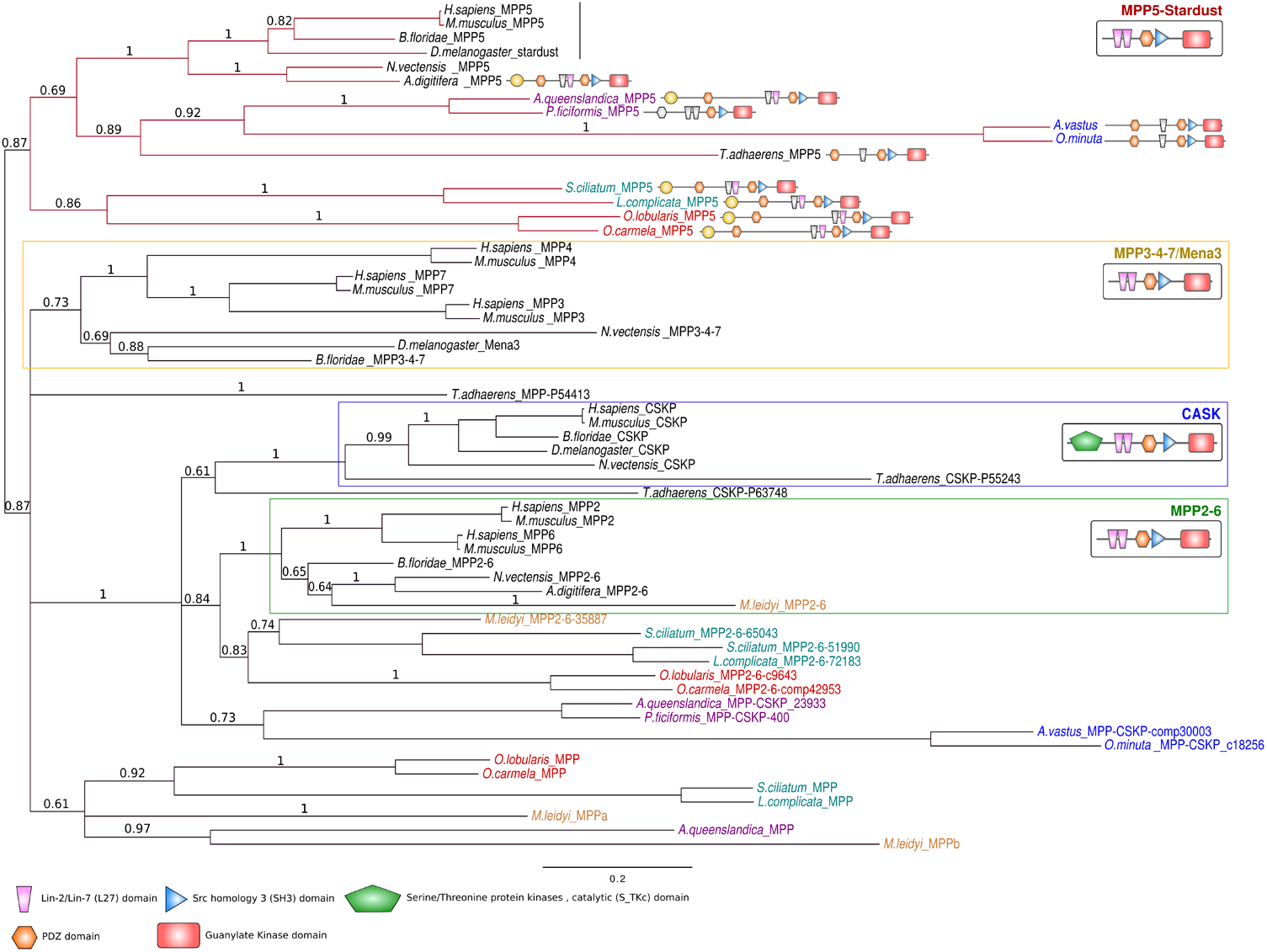
**Phylogenetic tree of MAGUK proteins** based on their shared MPP PDZ+SH3+GUK domains. A bayesian tree was inferred with available bilaterian sequences and predicted cnidarian, poriferan, placozoan and ctenophoran sequences aligned with MAFFT v7.123b. MrBayes was run under LG+G model of evolution with 4 rate categories and 1 million generations sampled every 1000 generations. The tree was rooted at midpoint and posterior probabilities are indicated for each branch. The canonical domain architecture was depicted for each main MPP class: MPP5-stardust, CASK, MPP2-6, MPP3-4-7. Whereas Ctenophora lack a MPP5/Sdt ortholog, all 4 sponge classes and Placozoa have one copy of the corresponding gene. Nevertheless, the first Lin2/Lin7 (L27) domain involved in interaction with the L27 domain of PatJ (Pals1-associated tight junction protein) is missing in glass sponges. Domains in grey are predicted but divergent.

## Discussion

By investigating the presence of genes and proteins involved in epithelial polarity and adherens junctions, we found that while Placozoa and Porifera (despite some divergence observed in hexactinellids) possess all polarity complex members and adherens junction components, ctenophores lack the Crumbs complex and Scribble. Indeed, we failed to find homologs of the corresponding genes in species available so far. In addition, *M. leidyi* possesses an E-cadherin-like cytoplasmic sequence divergent enough from canonical E-cadherin to raise doubt on its ability to interact with p120 catenin and with β-catenin. These unexpected findings are shedding a new light on the ongoing controversy about the morpho-anatomy of the last metazoan ancestor based on various phylogenetic reconstructions ^27,34,35,60–62^. One hypothesis favors ctenophores as a sister group of all other metazoans ^35,62^, suggesting that complex traits such as neurons and muscles might have been acquired independently in Ctenophora and Parahoxozoa ^60^. The alternative hypothesis favors Porifera as the sister group of other metazoans ^34,61^ in agreement with more traditional interpretations. According to our results, sponges (in particular Homoscleromorphs) now appear to have an epithelial toolkit (collagen IV, polarity complexes, E-cadherin complex) that is more complete (and expected to be functional according to motif conservation) than that of ctenophores. This finding is all the more unexpected since the epithelial organization of ctenophores appears fully admitted ^51^, while the presence of “true” epithelia in sponges remains debated.

In the context of the ctenophore-first evolutionary scenario, the compromised interactions between catenins and cadherins and the absence of two typical polarity complexes in ctenophora epithelia can be interpreted in two ways. Either it is the result of secondary losses, or it was inherited from an ancestral state. In this later case, it implies that additional components such as the Crumbs complex and p120 binding for E-cadherin were later acquired during the course of evolution in the last common ancestor of sponges and parahoxozoa.

Interestingly, the epithelial features of Ctenophora are very different ^14,26,27,63,64^ leading to incongruent interpretations of authors concerning the presence or not of bona fide AJs. According to our survey, if there are adhesive-type junctions in ctenophores, they cannot be considered as bilaterian AJ homologs.

Our results also challenge the notion that there is a straightforward relationship between a genetic toolkit and morphological features. Sponges from different classes possess very different tissue organizations; homoscleromorphs have epithelial-like layers with adherent junction while hexactinellids exhibit a syncitial organization without AJ-like junctions. Up to now, however, the small molecular variations observed in these different species are not sufficient to explain the huge differences seen in body plans. Similarly, for ctenophores, *Beroe*, *Pleurobrachia* and *Mnemiopsis* exhibit different epithelial features ^14^despite their very similar gene contents. Consequently, gene inventories alone are not sufficient to explain tissue and structure diversity.

Our study strongly advocates for more functional studies of the epithelium-like tissue of all non bilaterian animals. It is also an incentive to develop sponges and ctenophores as new experimental models for cellular biology to elucidate how the cell layer key to the rise of animal diversity emerged throughout evolution.

## Methods

### Genome sequencing and assembly

*Oscarella lobularis* genome sequencing was performed using Illumina technology with DNA-seq paired-end and Nextera mate pair protocols on a HiSeq2500 sequencer. Adapter sequences were removed and low-quality bases were trimmed using Cutadapt ^65^. Remaining reads were assembled using a pipeline including IDBA-UD (^66^, Platanus ^67^, GapFiller ^68^ and cap3 ^69^. A transcriptomic dataset was mapped with Tophat ^70^to all contigs longer than 1kb to identify potential Eukaryotic sequences. The result was passed to Braker ^71^to predict genes. All predicted protein sequences were submitted to BLAST (^72^to refine taxonomic assignment and automatically assess genes function. 17,885 protein-coding were predicted from a total of 2,658 contigs (Table S3). Following the mapping of the individual reads to the final genome and transcriptome assembly using Bowtie2 ^73^, the coverage values were found to be 98, ensuring a near 100% completeness of the predicted gene content.

*Oopsacas minuta* genome sequencing was first performed in the same conditions as *Oscarella lobularis*. In addition, a second genome sequencing step was performed using PacBio technology on an isolated sponge fragment to limit bacterial contaminations. These long reads were filtered based on their length and quality with Pacific Biosciences(PacBio)tools (SMRT Portal) then selfcorrected with canu ^74^. All Illumina reads were mapped on the corrected PacBio reads with Bowtie2. Mapped Illumina reads and corrected PacBio reads where then assembled together with SPAdes ^75^. The number of contigs longer than 1kb was low enough to rapidly identify Eukaryotic sequences using MetaGenemark ^76^and BLAST through a homebrew web server. Finally, a super scaffolding and polishing step was achieved using Sspace ^77^, Pilon ^78^and GapFiller. 17,043 genes were predicted with Braker from the 365 remaining contigs (see Table S3 for metrics). The same method as for *Oscarella lobularis* genome was applied to predict proteins and their functions. The mapping of the individual read to the Genome and transcriptome assembly resulted into an estimated coverage of 381, again ensuring a near 100% completeness of the gene content.

### Sequence annotation and structure prediction

Epithelial hallmarks were investigated using data from various sources: *O. lobularis* and *O. minuta de novo* assembled genomes and transcriptomes (this work), and on available poriferan, placozoan, cnidarian and ctenophoran genomes and/or transcriptomes retrieved from the sources listed in Table S4.*D. melanogaster*, *M. musculus* and *A. queenslandica* Epithelial cadherin, Crumbs, PAR and Scribble complexes retrieved from NCBI database were used to perform reciprocal besthits with BLAST 2.3.0 run locally using an E-value cutoff of 10^-5^.

*Ab initio* protein-coding gene prediction was performed on the best candidate genomic and/or transcriptomic contigs with GeneMark.hmm eukaryotic web serveur (http://exon.gatech.edu/gmhmme.cgi) ^79^, GenScan (http://genes.mit.edu/GENSCAN.html) ^80^, Augustus v3.0.3 (http://bioinf.uni-greifswald.de/augustus/) ^81^run locally and FgeneSH web server (http://www.softberry.com/) ^82^. Protein domains were predicted and checked with Pfam v28.0 (http://pfam.xfam.org/) run locally^83^, InterProScan v52 (http://www.ebi.ac.uk/interpro/) ^84^and SMART (http://smart.embl-heidelberg.de/) ^85^.

The newly identified early branching metazoan proteins were added to the previous database sequences for iterative BLAST searches to identify more potential homologs. Proteins containing repeated domains generate false positive and best-hits characteristic of protein motifs and /or domains were used to enhance the detection of real homologs. These motifs and domains were aligned with MAFFT.7 ^86^(http://mafft.cbrc.jp/alignment/server/) and/or MUSCLE v3.8.31 ^87^(implemented in Seaview 4.5.2 ^88^) depending on the level of conservation of the proteins. HMM profiles were built with HMMER 3.1b1 (http://hmmer.org/) using aligned sequences with an Evalue cutoff of 10^−5^. Retrieved motifs of potential homologs to the protein queries were used as new baits to recompute HMM profiles and perform further HMM searches. C-terminal characteristic parts were used to build HMM profiles of Crumbs and E-cadherin proteins.

Since Lin2/Lin7 (L27) (N)-terminal domain is important in the mediation of MAGUK family protein interactions with other proteins, a particular attention was given to this domain. To identify all potential MAGUK proteins, local BLAST searches (BLASTp and tBLASTn) were performed against complete or on-going genome projects (Table S4) using L27 domains retrieved from the Pfam database (PF02828). To improve the detection, L27 domain HMM profile was rebuilt by iteratively adding the best scoring L27 domains identified in sponges, placozoans, cnidarians and ctenophorans.

L27 domain HMM searches led also to the identification of PATJ homologs but for some proteins containing multiple PDZ domains (MPDZ) no L27 domain was detected. For MPDZ lacking the L27 domain, a BLASTp was run locally using the PDZ domains retrieved from the Pfam database (PF00595) as reference. This step led to the identification of *Petrosia ficiformis* MPDZ which has (N)- and (C)-terminal regions predicted on two different contigs, including L27 and PDZ domains on the first contig and the remaining PDZ domains on a second contig merged with Emboss merger webtool (http://www.bioinformatics.nl/cgi-bin/emboss/merger).

In addition to protein domain analyses, conservation of critical residues was investigated on aligned sequences according to previous publications. Alignments were visualized with JalView 2.9 using Clustalx amino acids color display ^89^.

### Phylogenetic analyses

To confirm the annotation of identified sponge genes as well as other early branching metazoans, a maximum likelihood (ML) analysis with 100 bootstraps was conducted using PhyML v3.1 ^90^(implemented in Seaview 4.5.2). Bayesian analyses were conducted using MrBayes v3.2.5 ^91^.Both ML and Bayesian analyses were run under the appropriate model recommended by ProtTest v3.4 ^92^.

For PALS1/MPP5/Stardust phylogeny, a first step consisted on inferring an ML and a Bayesian tree from SH3+GUK domains of the “core MAGUK” of all proteins containing GUK domains retrieved from NCBI or predicted sequences of available non-bilaterian genome and/or transcriptomes (data not shown). This phylogeny includes MAGUK, Dlg, LRR and GUK domain, Zonula occludens (ZO), Caspase recruitment family (CARMA). Membrane-associated Guanylate kinase Inverted (MAGI) and Calcium channel β-subunit (CACNB) classes were also included even MAGI do not have a SH3 domain, their GUK domain is truncated, and Calcium Voltage-gated Channel auxiliary subunit beta (CACNB) are divergent.

Once all proteins of MPP classes were clearly identified, another phylogenetic analysis was performed including the PDZ domains besides the SH3+GUK domains previously used.

The PATJ/MUPP-1/DLG/LIN phylogeny was performed on sequences identified using the L27 domains and the two following PDZ domains (except for the LIN family that exhibits a single PDZ domain).

All sequences annotated and analyzed are listed in Tables S5 and S6.

## ACKNOWLEDGMENTS

We thank Chris Toret for critical reading of this manuscript, the diving service of Observatoire des Sciences de l’Univers (Pythéas institute) for collecting *O. lobularis and O. minuta* samples and the Service Commun de Biologie Moléculaire (SCBM) of IMBE for providing facilities needed to prepare samples for sequencing.This work was supported by a grant from A*MIDEX (n° ANR-11-IDEX-0001-02)funded by the “investissements d’Avenir” French Government program, managed by the French National Research Agency (ANR). We acknowledge the use of the PACA-Bioinfo Platform, supported by IBISA and France-Génomique (ANR-10-INBS-0009). The Le Bivic group is an “Equipe labellisée 2008 de La Ligue Nationale contre le Cancer” and is supported by the labex INFORM (grant ANR-11-LABX-0054), CNRS and Aix-Marseille University.

## CONTRIBUTIONS

HB: sequences retrieval and analysis and manuscript writing ER:project design, data analysis and manuscript writing SS:genome and transcriptome analysis, manuscript editing CJ:genome and transcriptome analysis, manuscript editing JMC:project design and manuscript writing CB: project design, data CB: project design, data analysis and manuscript writing ALB:project design, data analysis and manuscript writing

---

**Figure S1A.**
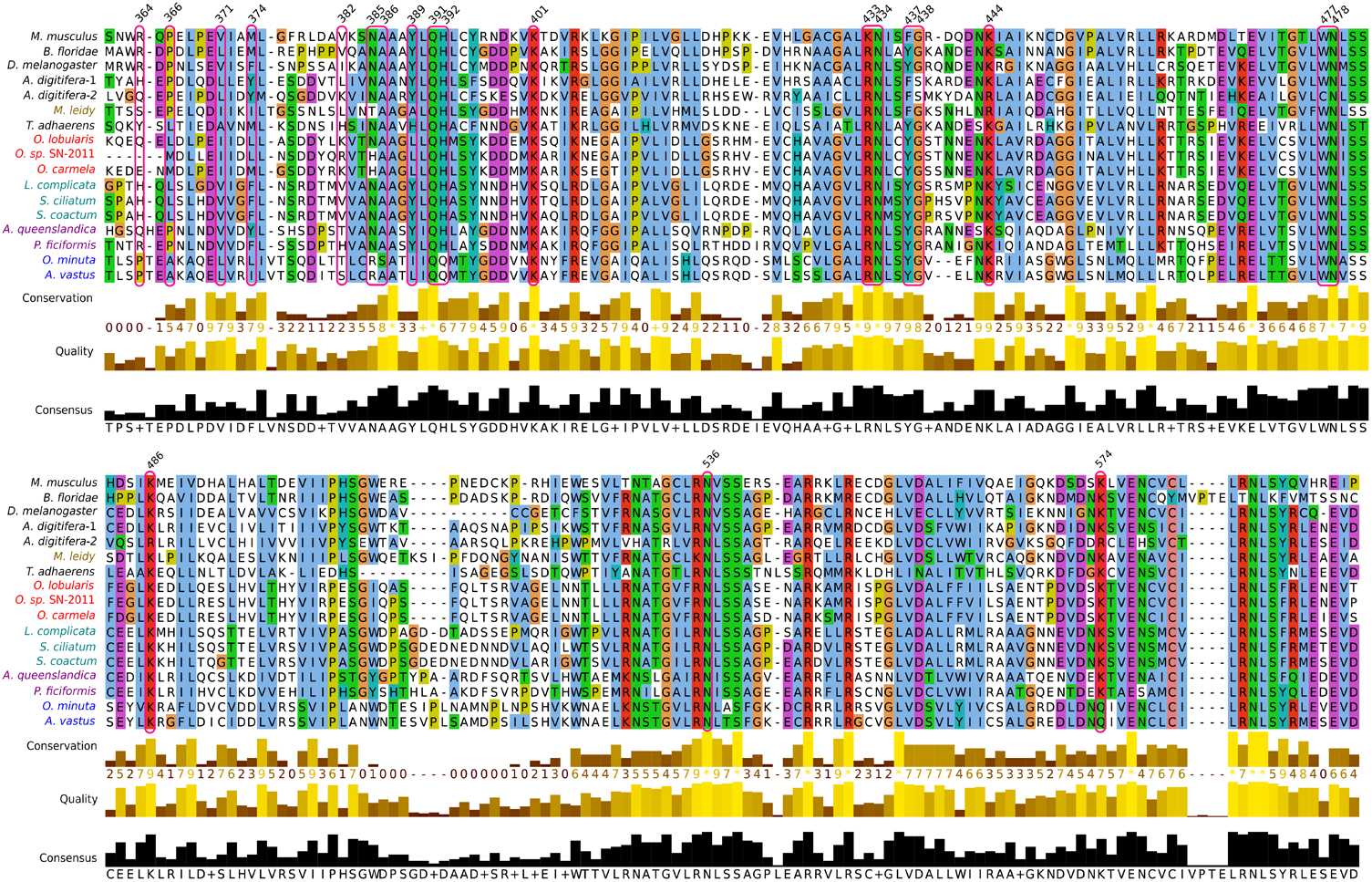
Comparison of p120 sequences. Residues involved in interaction with E-cadherin are boxed in red. Most of them are conserved.

**Figure S1B.**
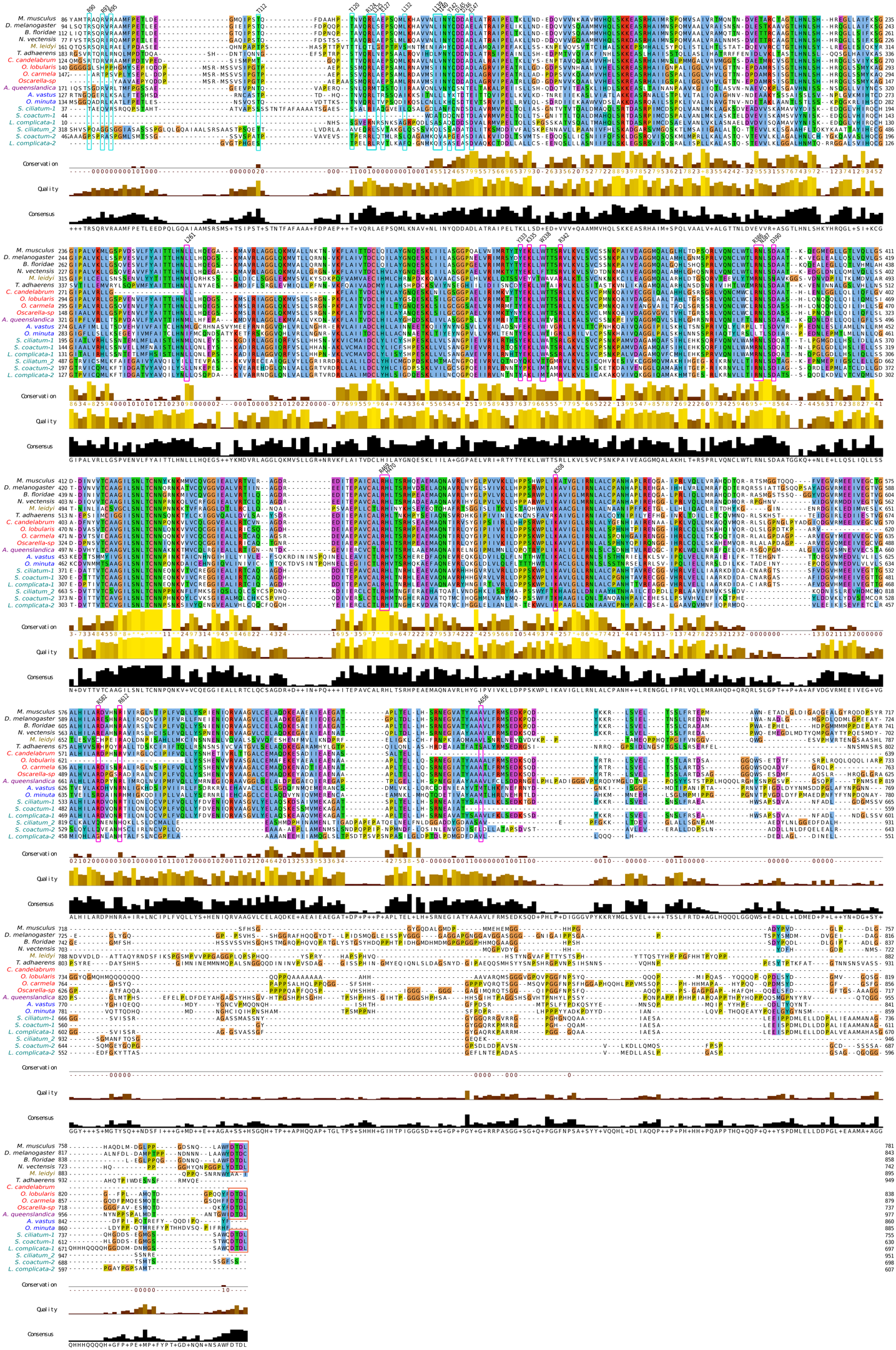
Comparison of β-catenin sequences. A single β-catenin gene copy was identified in every studied species except for calcareous sponges that exhibit a duplication. All residues essential for E-cadherin interaction are boxed in pink and are highly conserved except for the R386 and N387 residues (replaced by L and T, respectively) in two hexactinellids and a more anecdotal change from A656 to S in placozoans. Residues boxed in blue are involved in α-catenin binding and in orange for the DTDL PDZ binding motif.

**Figure S1C.**
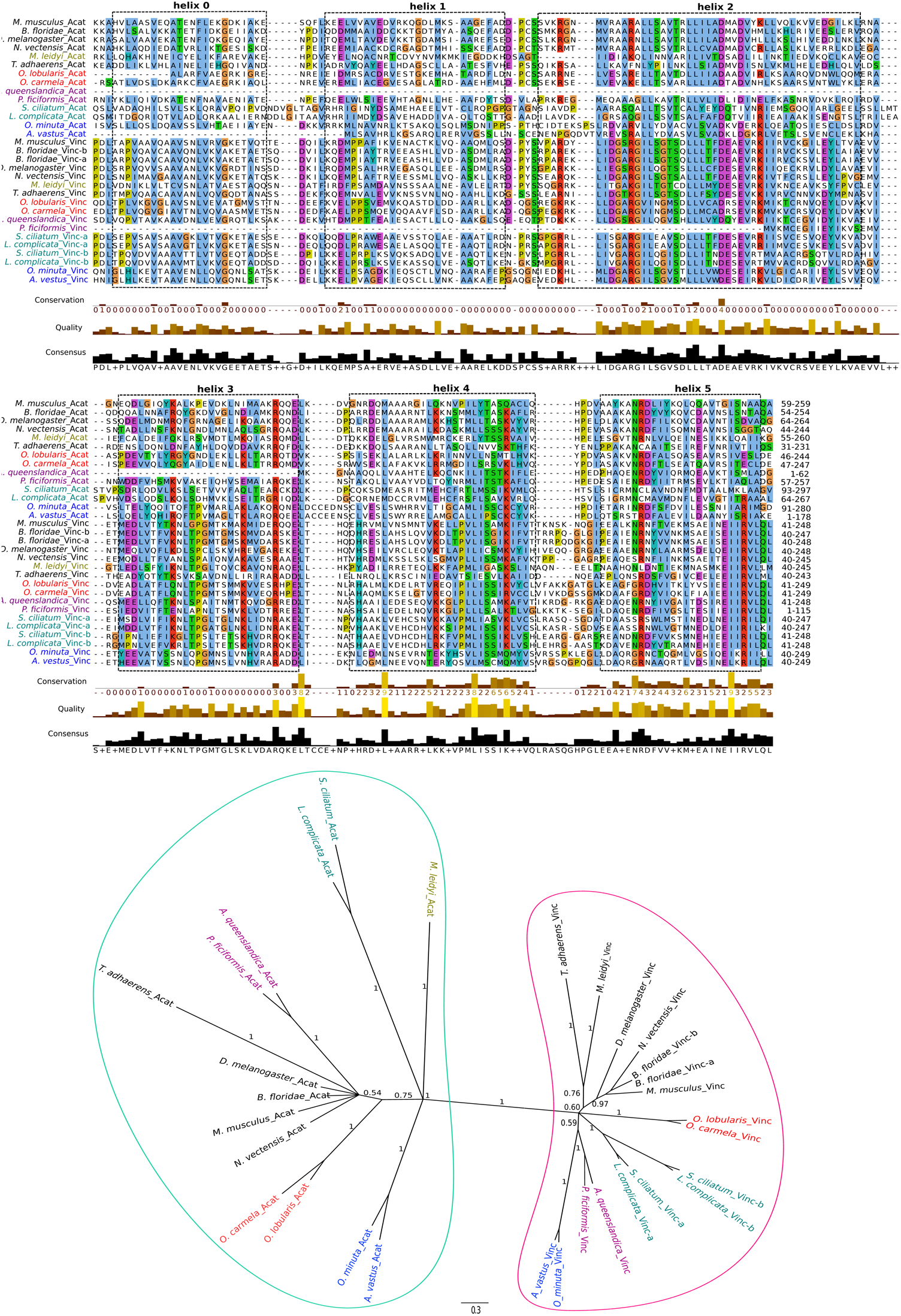
Analyses of α-catenins and vinculins sequences. Sequences of α-catenins and vinculins were aligned based on the structural domains helix0 to helix5 in *Mus musculus* α-catenin and vinculin. Helices are boxed and the numbers at the end of each sequence indicate the range encompassed in the alignment. Secondary structure prediction by JNet (Jalview option) identified six helices in all sponge α-catenin sequences except for *A. queenslandica* (missing the 4 first helices) and *A. vastus* (missing helix0). All species analyzed in this study have one copy of α-catenin and one copy of vinculin well-separated in Bayesian tree with high support (pp=1) (bottom).

**Figure S2.**
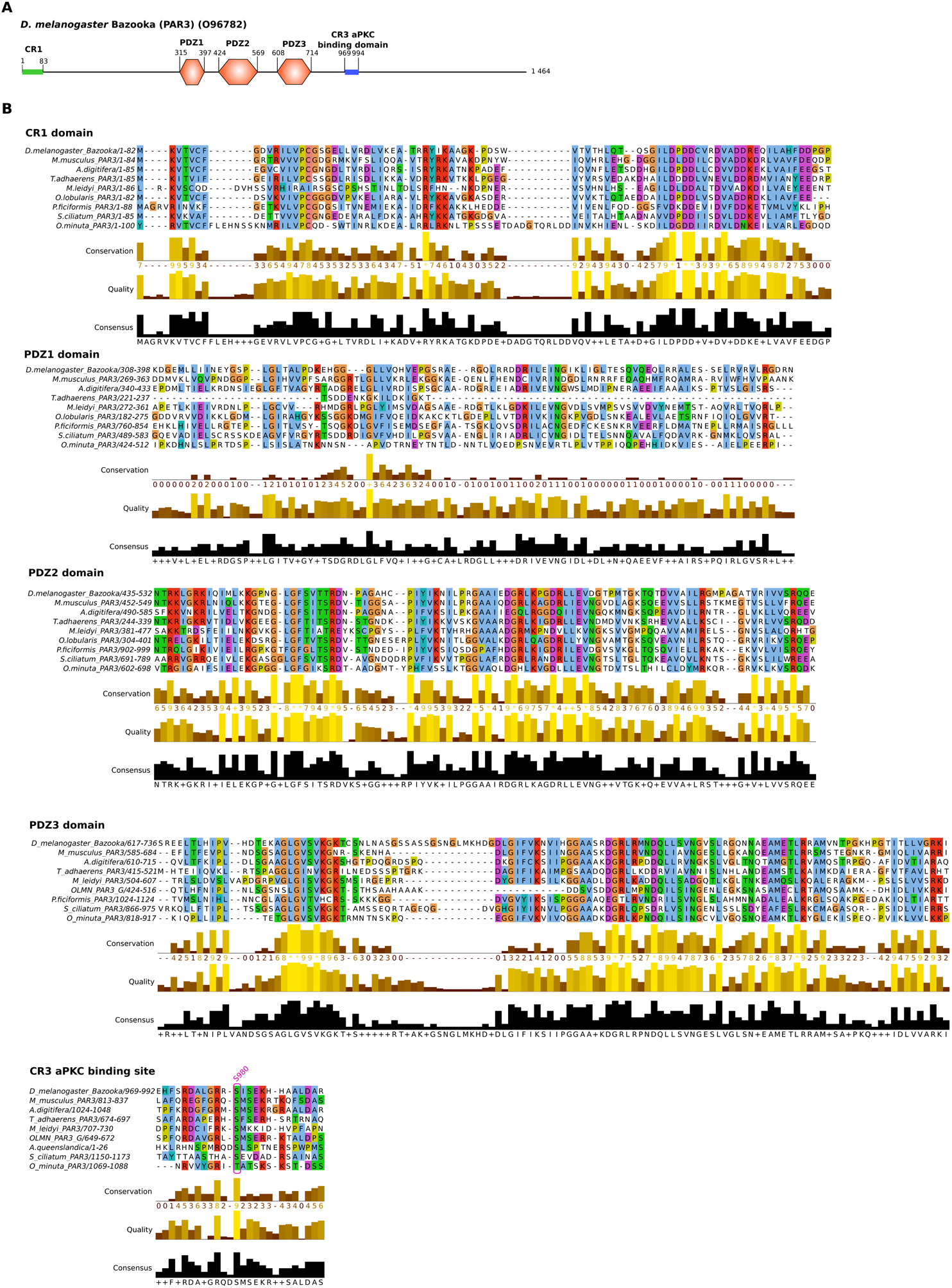
Structure of Par3 proteins in metazoans. Par3 exhibits a conserved N-terminal domain (CR1), three central PDZ domains, and a C-terminal region containing multiple protein binding sites including the aPKC-binding motif.

**Figure S5.**
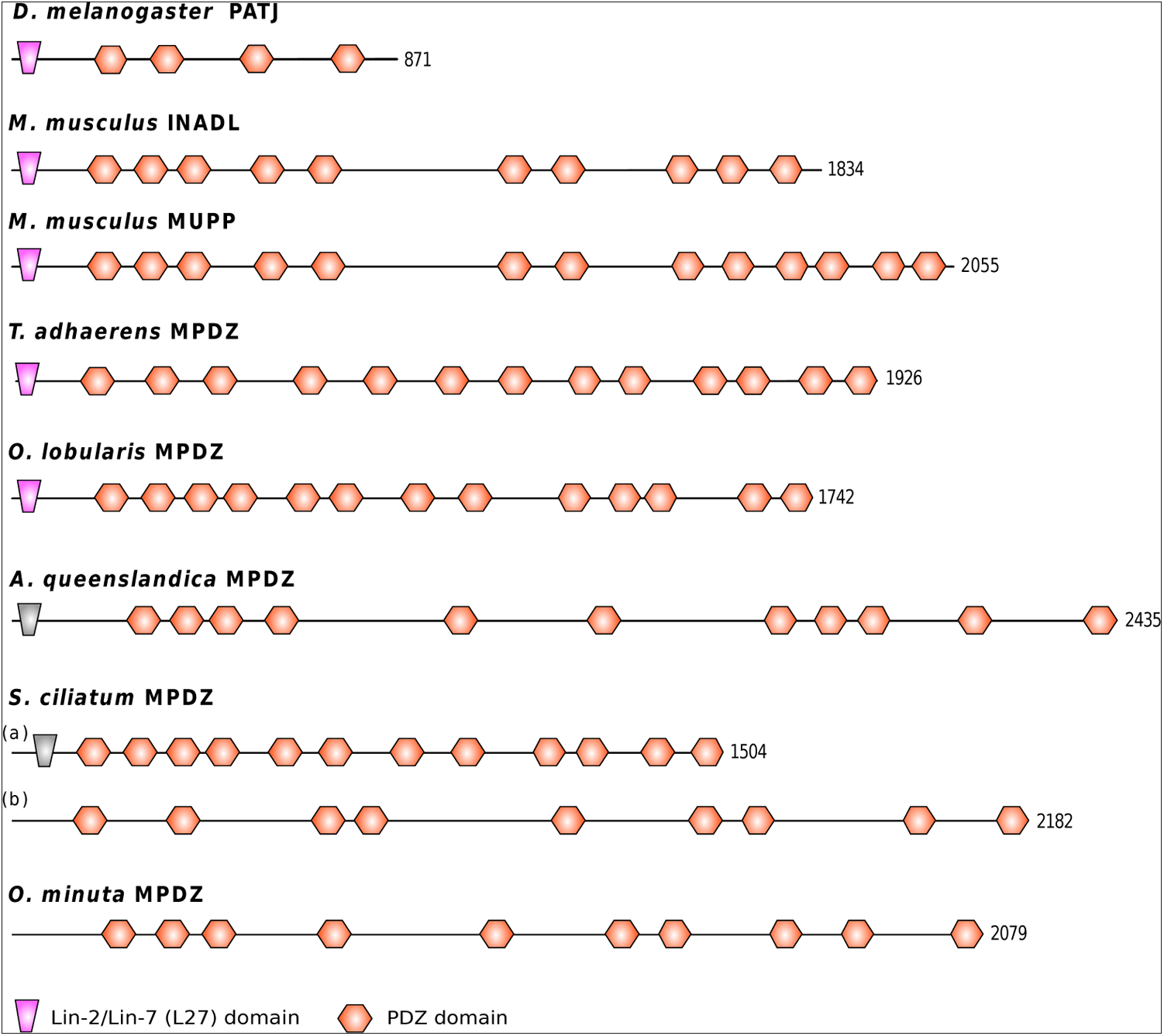
Domain composition of PatJ. (*D. melanogaster*), INADL and MUPP1 (*M. musculus*) and Multiple PDZ containing protein (MPDZ) (*O*. *lobularis*, *S. ciliatum*, *A. queenslandica* and *O. minuta*). Note that only *O. lobularis* exhibits an MPDZ with a well-detected L27 domain (Evalue=8.5 10^-4^) as bilaterians. *A. queenslandica and S. ciliatum* MPDZ have a low-scoring L27 domain (shaded in grey) according to the HMM profile search. There is no recognizable similarity to the L27 domain in the N-terminal region *of O. minuta* MPDZ.

**Table S1.**
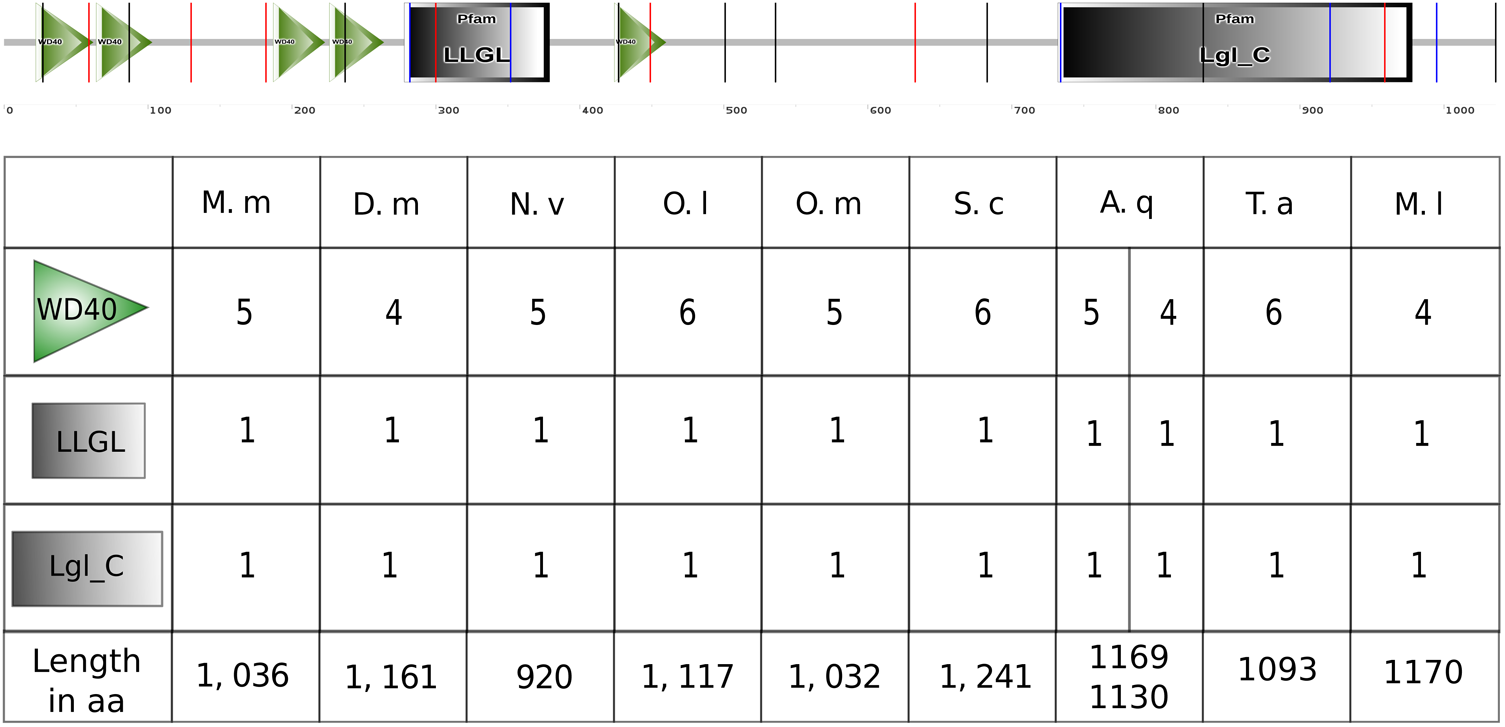
**Domain structures of the Lethal giant larvae (LGL) proteins in various metazoans** SMART domain diagram of *M. musculus* LGL1 (Q80Y17) canonical domain structure, and number of protein domains WD40, LLGL: Lethal giant larvae homologue 2 (Interpro domain IPR013577) and Lgl_C: Lethal giant larvae(Lgl) like, C-terminal (Pfam PF08596) for *M. musculus* (M. m); *Drosophila melanogaster* (D. m); *Nematostella vectensis* (N. v); *Oscarella lobularis* (O. l); *Oopsacas minuta* (O. m); *Sycon ciliatum* (S. c); *Amphimedon queenslandica* (A. q); *Trichoplax adhaerens* (T. a) and *Mnemiopsis leidyi* (M. l)

|  | M. m | D. m | N. v | O. l | O. m | S. c | A. q |  | T. a | M. l |
| --- | --- | --- | --- | --- | --- | --- | --- | --- | --- | --- |
|  | 5 | 4 | 5 | 6 | 5 | 6 | 5 | 4 | 6 | 4 |
|  | 1 | 1 | 1 | 1 | 1 | 1 | 1 | 1 | 1 | 1 |
|  | 1 | 1 | 1 | 1 | 1 | 1 | 1 | 1 | 1 | 1 |
| Length in aa | 1, 036 | 1, 161 | 920 | 1, 117 | 1, 032 | 1, 241 | 1169 | 1130 | 1093 | 1170 |

**Table S2.**
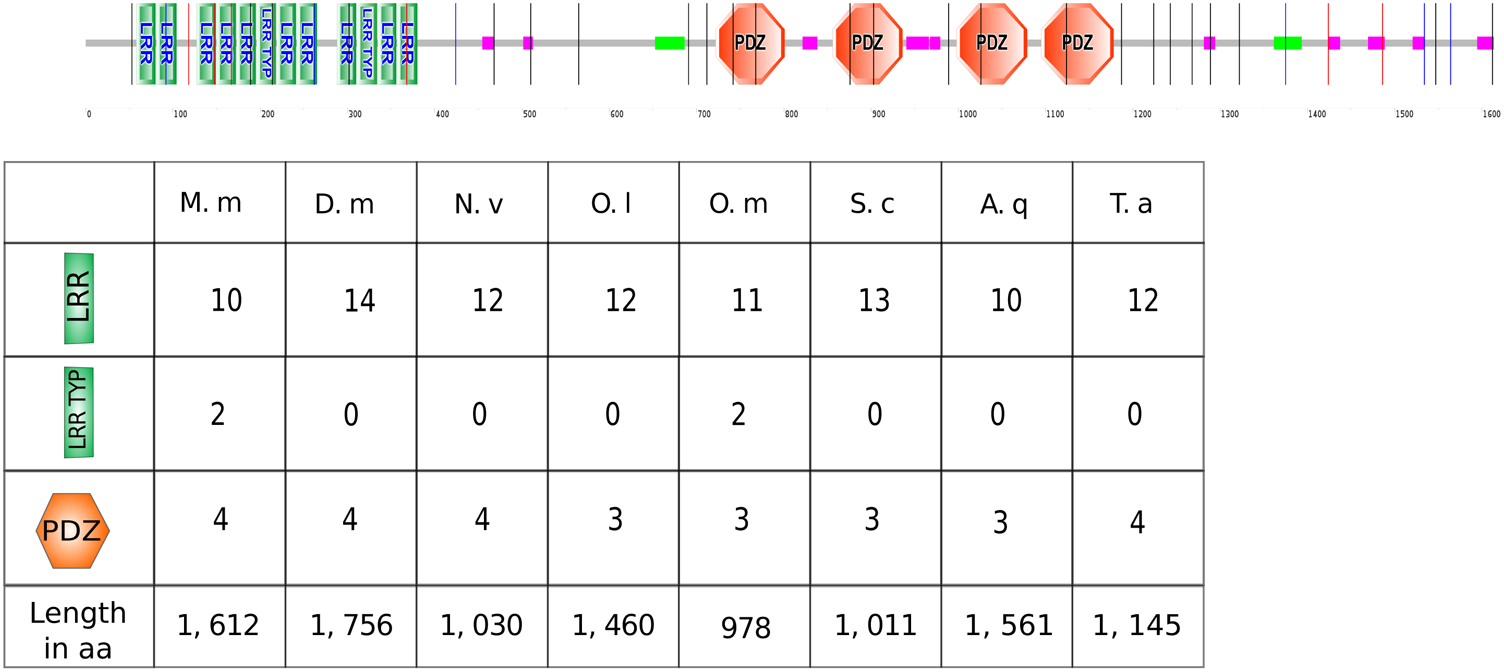
**Domain structures of the Scribble (Src) proteins in various metazoans** SMART domain diagram of *M. musculus* Scribble (Q80U72) canonical domain structure, and number of protein domains LRR (leucine rich repeats), LRR TYP (leucine rich repeats, typical) and PDZ for *M. musculus* (M. m); *Drosophila melanogaster* (D. m); *Nematostella vectensis* (N. v); *Oscarella lobularis* (O. l); *Oopsacas minuta* (O. m); *Sycon ciliatum* (S. c); *Amphimedon queenslandica* (A. q); *Trichoplax adhaerens* (T. a) and *Mnemiopsis leidyi* (M. l). *M. leidyi* possesses multiple proteins containing LRR domains but since none PDZ domain is associated, these proteins were not taken into acoun

|  | M. m | D. m | N. v | O. l | O. m | S. c | A. q | T. a |
| --- | --- | --- | --- | --- | --- | --- | --- | --- |
|  | 10 | 14 | 12 | 12 | 11 | 13 | 10 | 12 |
|  | 2 | 0 | 0 | 0 | 2 | 0 | 0 | 0 |
|  | 4 | 4 | 4 | 3 | 3 | 3 | 3 | 4 |
| Length in aa | 1, 612 | 1, 756 | 1, 030 | 1, 460 | 978 | 1, 011 | 1, 561 | 1, 145 |

**Table S3:** Characteristics of the new private databases used in this study

|  | Taxa | species | Dataset | raw reads | corrected/selected reads | mapped reads | depth |
| --- | --- | --- | --- | --- | --- | --- | --- |
| Porifera | Hexactinellida |  | Pacbio | 751 460 | 124 793 | 112 130 |  |
|  |  | Leucopsacidae | PE <sup>1</sup> | 395 851 746 | 335 361 472 | 234 150 237 | 380 |
|  |  | *Oopsacas minuta* | MP<sup>2</sup>_3-5kb | 245 157 912 | 92 657 868 | 24 712 816 | 54 |
|  |  |  | MP_5-8kb | 274 964 868 | 116 755 522 | 25 643 839 | 56 |
|  |  |  | MP_8-11kb | 191 742 544 | 79 280 536 | 15 423 954 | 34 |
|  |  |  | Transcriptome | 421 126 244 |  | 207 529 788 | 1 443 |
|  | Homoscleromorpha |  | PE | 307 384 094 | 112 751 740 | 51 225 276 | 97 |
|  |  | Oscarellidae | MP_3-5kb | 254 563 152 | 106 866 702 | 20 327 965 | 51 |
|  |  | *Oscarella lobularis* | MP_5-8kb | 251 314 628 | 106 866 702 | 18 165 914 | 45 |
|  |  |  | MP_8-11kb | 295 517 160 | 114 050 246 | 13 598 285 | 34 |
|  |  |  | Transcriptome | 380 096 086 |  | 231 475 388 | 710 |

1: Paired end
2: Mate Pair

**Table S4:**
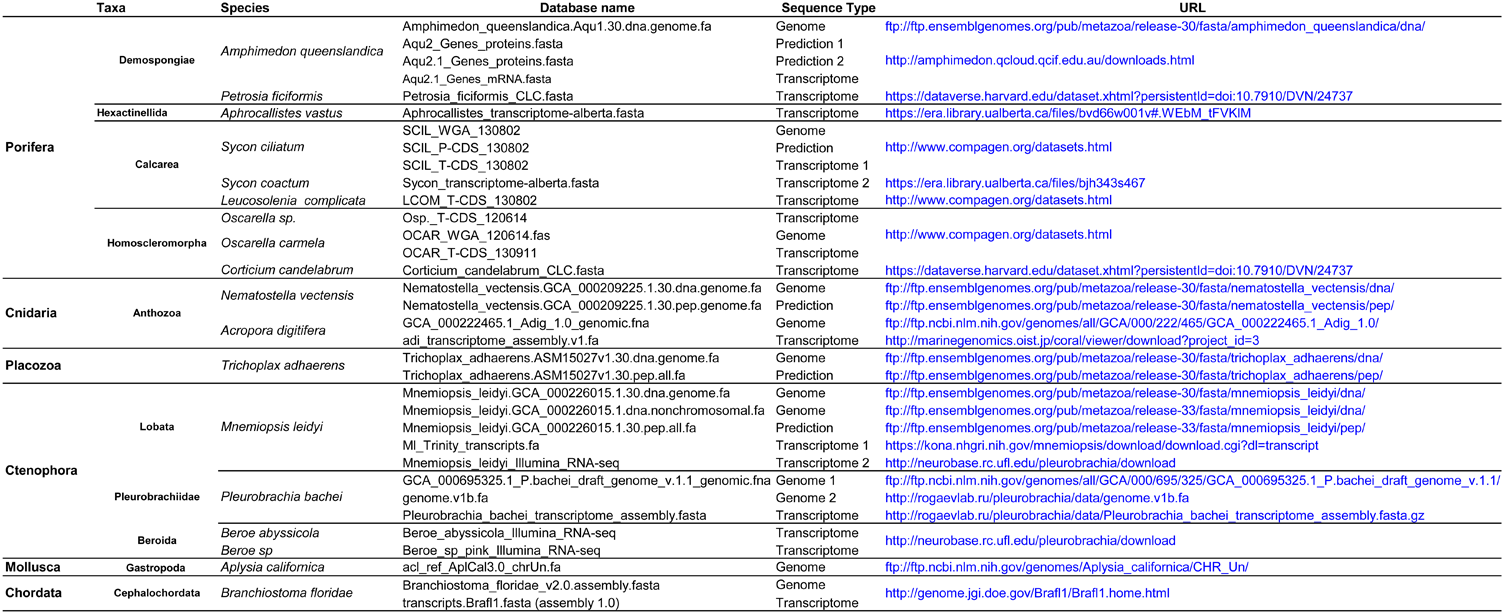
Public databases used in this study: nature (genome/transcriptome) and link for accession

**Table S5:**
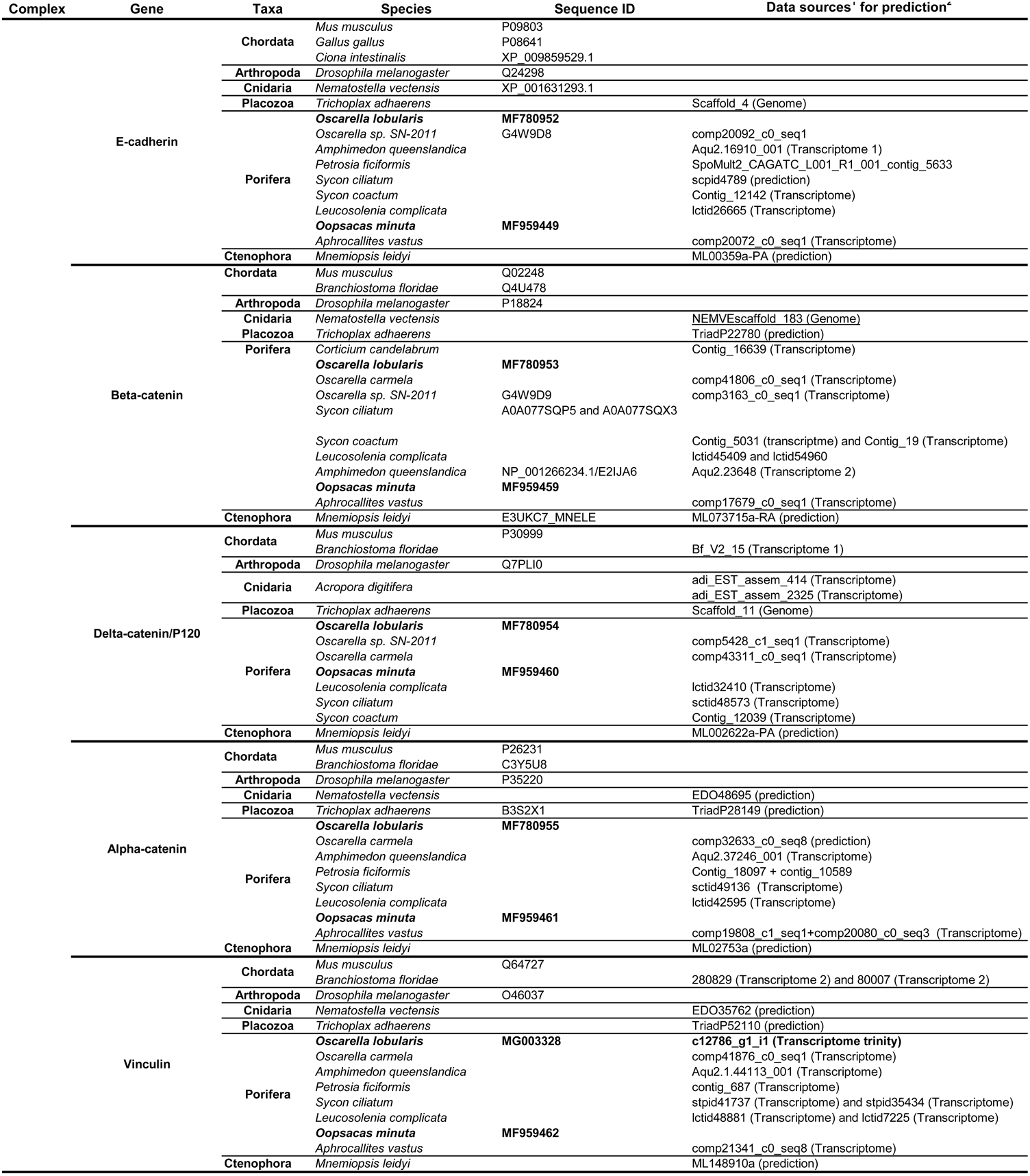
Cadherin-Catenin Complex (CCC) sequence retrieval: Accession numbers or contig/scaffold references where candidate genes were identified. In bold accession numbers of sequences annotated from our two new and genomic sponge datasets. 1: see table S4 for the links to corresponding data sources; 2: Prediction of the protein sequence and/or the function

**Table S6:**
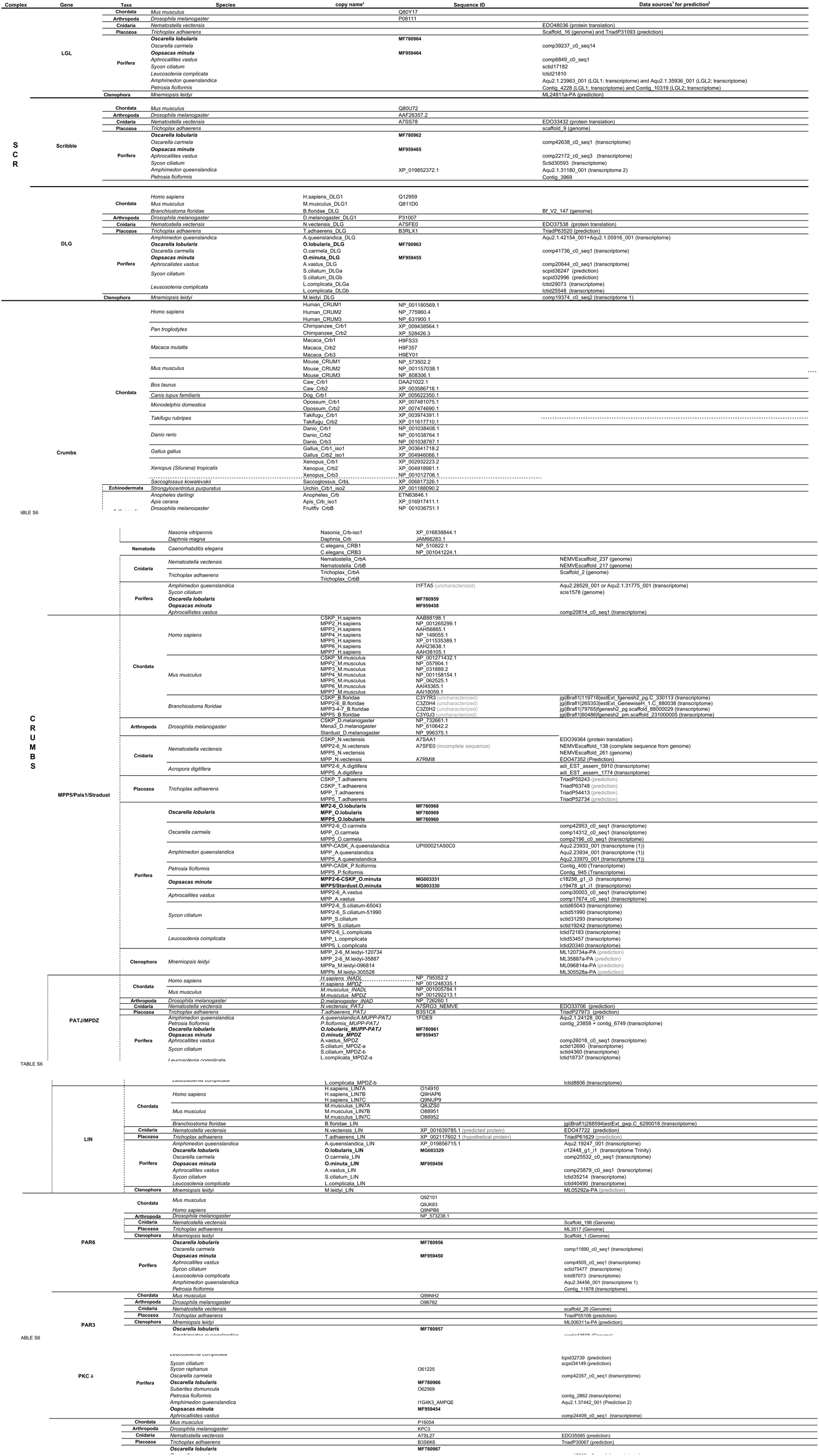
Polarity complex sequence retrivel: Accession number of contig/scaffold where candidate genes were identified. Sequences from our new genomic and transcriptomic datasets are in bold 1: links to corresponding data sources are provided in table S4; 2: Prediction of the protein sequence and/or the fuction 3:names used for multigenic families

